# Heterotypic Aβ interactions facilitate amyloid assembly and modify amyloid structure

**DOI:** 10.1101/2021.04.28.441786

**Authors:** Katerina Konstantoulea, Patricia Guerreiro, Meine Ramakers, Nikolaos Louros, Liam Aubrey, Bert Houben, Emiel Michiels, Matthias De Vleeschouwer, Yulia Lampi, Luís F. Ribeiro, Joris de Wit, Wei-Feng Xue, Joost Schymkowitz, Frederic Rousseau

## Abstract

It is still unclear why pathological amyloid deposition initiates in specific brain regions, nor why specific cells or tissues are more susceptible than others. Amyloid deposition is determined by the self-assembly of short protein segments called aggregation-prone regions (APRs) that favour cross-β structure. Here we investigated whether Aβ amyloid assembly can be modified by heterotypic interactions between Aβ APRs and short homologous segments in otherwise unrelated human proteins. We identified heterotypic interactions that accelerate Aβ assembly, modify fibril morphology and affect its pattern of deposition *in vitro*. Moreover, we found that co-expression of these proteins in an Aβ reporter cell line promotes Aβ amyloid aggregation. Importantly, reanalysis of proteomics data of Aβ plaques from AD patients revealed an enrichment in proteins that share homologous sequences to the Aβ APRs, suggesting heterotypic amyloid interactions may occur in patients. Strikingly, we did not find such a bias in plaques from overexpression models in mouse. Based on these data, we propose that heterotypic APR interactions may play a hitherto unrealised role in amyloid-deposition diseases.

## Introduction

Neurodegenerative amyloid diseases are a diverse group of pathologies that present very different symptoms and progressions in different areas of the brain (Chiti & Dobson, 2017; Goedert *et al*, 2017; Walsh & Selkoe, 2016). Simultaneously these diseases share common hallmarks that remain poorly explained. First, their initiation is characterized by the deposition of particular proteins in specific cells or brain regions (Gan *et al*, 2018). Second, this process is associated to functional and homeostatic dysregulation of affected cells ultimately resulting in neuronal death (Zaman *et al*, 2019). Third, from the site of initiation the disease progresses in a stereotypical manner by propagation of amyloid deposition to anatomically connected cells and brain regions with symptoms that match their function (Taylor *et al*, 2002). Together these properties suggest specific neuronal and regional vulnerability of the brain to the aggregation, propagation and toxicity of particular amyloidogenic proteins (Fu *et al*, 2018; Muratore *et al*, 2017). Factors enhancing these vulnerabilities include physiological ageing but also disease-specific familial mutations and population risk factors (Hipp *et al*, 2019; Silva *et al*, 2019). This illustrates how neuronal susceptibility is favoured by general (protein) homeostatic ageing but that disease initiation and its effects are highly context dependent. It is still unclear which cellular interactions contribute to the modulation of neuronal susceptibility either by sensitizing or protecting particular neurons or brain regions to aggregation. It is also not known whether amyloid interactions in each of these diseases are purely idiosyncratic or whether cross-β amyloid structure also favours canonical modes of interaction that provide generic mechanisms for amyloid gain-of-function.

Amyloid structures from different proteins grown either in vitro or in vivo share a common cross-β architecture (Gallardo *et al*, 2020; Landreh *et al*, 2016; Lutter *et al*, 2019; Riek & Eisenberg, 2016). Structural analysis of disease-associated amyloid structures and their polymorphs revealed that they are not uniformly stable but that some regions dominate the thermodynamic stability of the amyloid (van der Kant *et al*, 2021). Interestingly stable regions correspond to those previously identified as the amyloid nucleating segments of these proteins (Ganesan *et al*, 2016; Marshall *et al*, 2016; Teng & Eisenberg, 2009; Ventura *et al*, 2004). These aggregation-prone regions (APRs) consist of short sequence segments, 5-15 residues in length (Fernandez-Escamilla *et al*, 2004b; Goldschmidt *et al*, 2010; Rousseau *et al*, 2006b) and their thermodynamic stability results from their high propensity to adopt the cross-β conformation (Louros *et al*, 2020; Rousseau *et al*, 2006a; van der Kant *et al.*, 2021). APRs assemble into stable amyloids both on their own as peptides or in the context of full-length proteins underlining their essential (Ganesan *et al.*, 2016; Marshall *et al.*, 2016) and dominant role (Teng & Eisenberg, 2009; Ventura *et al.*, 2004). Because of these favourable conformational properties, APRs constitute good protein interaction interfaces favouring amyloid self-assembly, i.e. through their affinity for binding to their own sequence (Krebs *et al*, 2004; O’Nuallain *et al*, 2005; O’Nuallain *et al*, 2004; Vanik *et al*, 2004; Wetzel, 2006). Recent evidence however suggests that amyloid self-specificity is not absolute and that disease amyloids can engage heterotypic interactions resulting in cross-seeding and co-aggregation (Giasson *et al*, 2003; Konstantoulea *et al*, 2021; Lutter *et al.*, 2019; Ly *et al*, 2021; Oskarsson *et al*, 2015) that are relevant to the pathophysiology of these disease (Colom-Cadena *et al*, 2013; Gallardo *et al.*, 2020; Pham *et al*, 2019; Sampson *et al*, 2020; Vasconcelos *et al*, 2016). The fact that sequence similarity is apparent in many of these cross-interactions (Konstantoulea *et al.*, 2021) - and especially with APRs - suggests that APRs constitute favoured protein interaction interfaces for heterotypic protein interactions.

Here we investigated the potential of amyloid core APRs to engage in heterotypic amyloid interactions with human proteins that share local sequence homology with amyloid APRs. Next, we evaluated the potential of such interactions to modify the structure and kinetics of assembly of amyloids. In order to do this, we used the Alzheimer beta peptide Aβ1-42 as a paradigm as it is a relatively short amyloid peptide sequence the kinetics of which are well-documented. Aβ harbors two APRs: APR1 encompassing residues (16-21) where several familial AD mutations cluster and APR2 at the C-terminal region (29-end), whose variable length is an important factor in the development of sporadic AD (Fernandez-Escamilla *et al.*, 2004b; Marshall *et al.*, 2016; Vandersteen *et al*, 2012). Both regions have a high aggregation propensity due to a high hydrophobicity and beta-sheet propensity (Fernandez-Escamilla *et al*, 2004a) and readily form amyloid-like aggregates by themselves as peptides (de la Paz & Serrano, 2004). The importance of these APRs for the amyloid formation of Aβ was further demonstrated by a variant form of Aβ1-42 that was designed to suppress both APRs by introducing a single amino acid substitution in each region (Marshall *et al.*, 2016). This variant, carrying two mutations in total, was shown to no longer aggregate, which also rescued the neurotoxicity of Aβ (Marshall *et al.*, 2016), showing that both APRs are indeed key determinants of the kinetics of amyloid formation of Aβ.

We identified several peptides with homology to Aβ1-42 APR derived from human proteins including proteins expressed in the brain and demonstrated that they are able to interact with Aβ1-42 and alter its aggregation kinetics and fibril morphology. Moreover, we showed that in the context of the full-length protein these same sequences favour Aβ1-42 aggregation in a reporter cell line. Importantly, reanalyzing deep proteomics data of human Aβ plaques (Xiong *et al*, 2019) we showed that proteins with homology to Aβ APRs are overrepresented in amyloid plaques from AD patients and that they cluster in gene ontologies related to synaptic organization and regulation of vesicle-mediated transport. An overrepresentation that is not seen in mouse APP overexpression models. Together our analysis demonstrates that at least in the case of Aβ amyloid assembly interfaces provided by APRs also allow for heterotypic interactions with other proteins by a mechanism of local sequence homology and that such interactions have the potential to modify amyloid nucleation, elongation and fibril morphology and co-opt such proteins into plaques.

## Results

### Nomenclature

Aβ: The Alzheimer beta-peptide, which exists as a mixture of different length due to carboxy- and amino-terminal heterogeneity resulting from its proteolytic generation.

Aβ1-42: A single form of the Aβ peptide, starting from canonical position 1 and ending in position 42. This species is particularly enriched in the plaques of patients with sporadic AD.

### Mapping Aβ self-interactions using peptide arrays

To set up a method to investigate self-interactions between Aβ molecules, we turned to a method that was previously successfully used for analysing self-interactions of the yeast prion Sup35 (Tessier & Lindquist, 2007), namely peptide arrays in which peptides correspond to a sliding window over a target protein. By exposing these arrays to an aggregation-prone protein, the location of self-interaction sites in the sequence can be observed directly, provided the interacting residues form a contiguous stretch as in the APR model. For this purpose, we synthesised in-house peptide arrays on a cellulose membrane, by using a sliding window scan over the sequence of Aβ1-42 of length 12 and step size 1 (Figure 1a, Appendix Table S1). Given that the most widely used antibodies against Aβ have linear epitopes and thus would show cross-binding on such arrays, we resorted to using a biotinylated derivative of Aβ1-42 (Biot-Aβ1-42, rPeptide) that avoids interference during detection. Upon dissolving Biot-Aβ1-42, we performed Size Exclusion Chromatography (SEC, S75, GE Healthcare) using an inline Multiple Angle Light Scattering detector (MALS, Wyatt) to determine the molecular size of the eluents. This revealed a monomeric peak eluting around 14 mL and an oligomeric peak, eluting in the void volume (Figure 1b). Then we exposed a peptide array to the monomeric fraction of Biot-Aβ1-42 (100 nM) for 1 hour and detected binding of Aβ to the spotted peptides using streptavidin-HRP, but we observed no significant binding to any of the peptide spots (Figure 1c). However, detection with the 4G8 (Chang *et al*, 2007), 6E10 and 12F4 antibodies that recognize a central 18-22 epitope in Aβ, the N-terminus and the C-terminus respectively showed clearly that these peptide sequences are present, confirming also the quality of synthesis (Appendix Figure S1). When we exposed a fresh membrane to the void volume fraction in the same way as the monomer, we observed clear binding to the N-terminal fraction of APR1 (Figure 1d), but not APR2. We left monomeric Biot-Aβ1-42 samples to aggregate (at 10 μM) while monitoring their aggregation kinetics using Thioflavin-T (ThT) fluorescence (Figure 1e). At three different time points during the course of the aggregation, we took samples incubated them in parallel without ThT and put them on a fresh peptide microarray (at 100 nM). Aggregating species collected during the lag phase of aggregation (sample 1 on Figure 1e) showed again a binding pattern in the amino-terminal peptides of APR1, similar to the void volume species observed during SEC-MALS (Figure 1f). Aggregating species, taken later in the kinetic from the early elongation phase (sample 2), showed binding throughout both APR1 and APR2 (Figure 1g). Finally, the predominantly fibrillar aggregates present during the plateau phase (sample 3), showed only very weak binding to the Aβ peptides (Figure 1h). Finally, we generated amyloid seeds in the reverse reaction, using 15cycles of 30s sonication to generate fragments from mature amyloid fibrils that were obtained after 14 days of incubation. We used the aggregation kinetics to confirm that the sonication protocol led to the formation of functional amyloid seeds, and indeed we found this sample produced a notable reduction of the lag-phase of aggregation of Biot-Aβ1-42 at 5% or 10% molar ratio in monomeric units (Figure 1i), whereas the mature fibrils did not have this effect. These ‘reverse seeds’ (i.e. generated from mature fibrils) indeed also showed an interaction pattern with the membrane that was similar to that of the late oligomers of the elongation phase, with interactions across both APR regions (Figure 1g).

**1.**
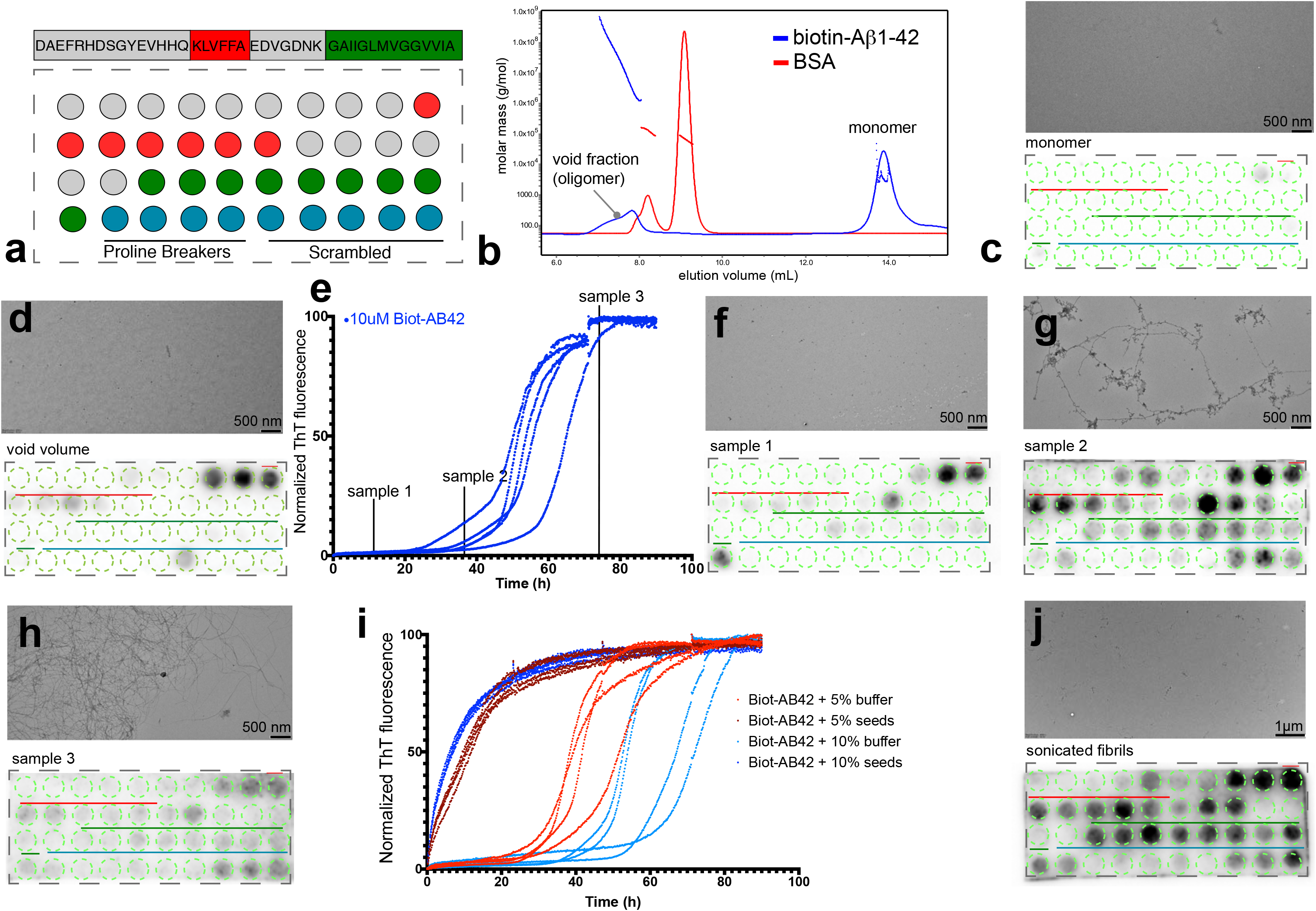
Differential binding of Aβ aggregating species in Aβ cellulose peptide microarrays. a) Aβ1-42 sliding window membrane setup. Red indicates where the KLVFFA starts presented whole. Green where GAIIGL presented whole. Blue indicates the controls (4 proline breakers, 5 scrambled Aβ peptides, sequences Supplementary Table1,3,5). b) SEC-MALS of Biot-Aβ1-42 preparation with 7M GnHCL showing a clear monomeric peak and a smaller oligomeric. c)100nM of Biot-Aβ1-42 monomers show low binding on membrane (down panel), TEM image shows no aggregating species in the sample (upper panel) d) Void fraction (oligomers) show strong binding on first APR of Aβ1-42. e) Normalized ThT kinetics of 10μM Biot-Aβ1-42 with timepoints of samples that incubated with Aβ1-42 membranes. f-h) Binding of different aggregating samples to Aβ membranes and their TEM images. 100nM of sample1 (early oligomers) binds strongly to middle APR (f), 100nM of sample2 (late oligomers) binds in both middle and C-terminal APR of Aβ1-42 (down panel) while TEM images shows fibrillar structures (upper panel) (g), 100nM of sample3 shows no specific binding to Aβ1-42 membranes(h). i) ThT kinetics of Biot-Aβ1-42 seeding. 10μM of Biot-Aβ1-42 incubated with 0.5 or 1 μM of Biot-Aβ1-42 seeds. j) 100nM of Biot-Aβ1-42 seeds show a strong binding in both APRs.

The difference in binding between monomeric and oligomeric Biot-Aβ1–42 species is consistent with the nucleation-growth kinetics of amyloid aggregation, in which the rate-limiting step is the formation of the oligomers, to which monomer addition is then relatively rapid (Dobson, 1999). Hence the preformed oligomers interact much more than the monomer, and the mature fibrils have few interaction sites. Interestingly, the peptide array data also shows that the early Biot-Aβ1–42 oligomeric intermediates (found in the fresh sample or formed from the monomer fraction) engage in different molecular interactions than later oligomeric species, which essentially behave as the fibril fragments generated by sonication. Consistent with the notion that the APRs are the kinetic determinants of Aβ aggregation, we indeed found the oligomers to interact mainly with peptides on the membranes corresponding to these regions. Sequences from APR1 from the central region of Aβ seems to form more interactions with early oligomers, whereas later oligomeric species also interacted with APR2 from the C-terminus.

### Protein fragments with local sequence similarity that bind to Aβ APRs occurs throughout the proteome

The APR regions of Aβ vary in length from 6 to more than 10 amino acids, a size distribution similar to that of APRs in other amyloids. Although most sequences of length 7 amino acids and longer are unique within the human proteome (Ganesan *et al*, 2015), we wondered how much sequence similarity exist within the proteome when considering mismatches. To this end, we plotted the number of similar sequence matches found in the human proteome (up to 2 mismatches) in function of fragment length, based on 1000 randomly selected human protein fragments per length, allowing up to 2 mismatches (Figure 2a) (proteome obtained from UniProt (UniProt, 2008), reviewed/unreviewed entries, filtered for 90% redundancy using CD-Hit (Fu *et al*, 2012)). This plot shows that the number of sequence matches drops exponentially with the length of the fragment and levels off at a length of 9 amino acid residues. Incidentally, this is the length that the immune system employs for self-discrimination, i.e. the length of the peptides displayed by the Major Histocompatibility complex (MHC) start at 9 amino acids. Given this strong dependence on fragment length of the number of homologous matches found for any given sequence in the proteome, we chose to fix this parameter in order to compare between APRs. Hence, we decided to use a relatively low fragment length of 6 to combine some specificity with a large candidate pool. Thus, we performed a search of the human proteome for hexapeptides matching KLVFFA and LVFFAE, allowing up to two mismatches, yielding 4390 matches, not filtering for isoforms (Appendix Table S2). We chose the middle region APR peptides since in our peptide microarrays showed strong binding with both early and late oligomers. Apart from Aβ and the parental amyloid precursor protein (APP), the only other proteins with an identical match are the Potassium voltage-gated channel subfamily B, members 1 and 2, which have a perfect match to LVFFAE towards the extracellular region of a transmembrane region. In addition, we identified 61 matches with a single mutation and 4318 matches with 2 mutations. The composition of the mismatches appears to follow a fairly random distribution (Figure 2b and c), with most amino acids appearing at each position.

**2.**
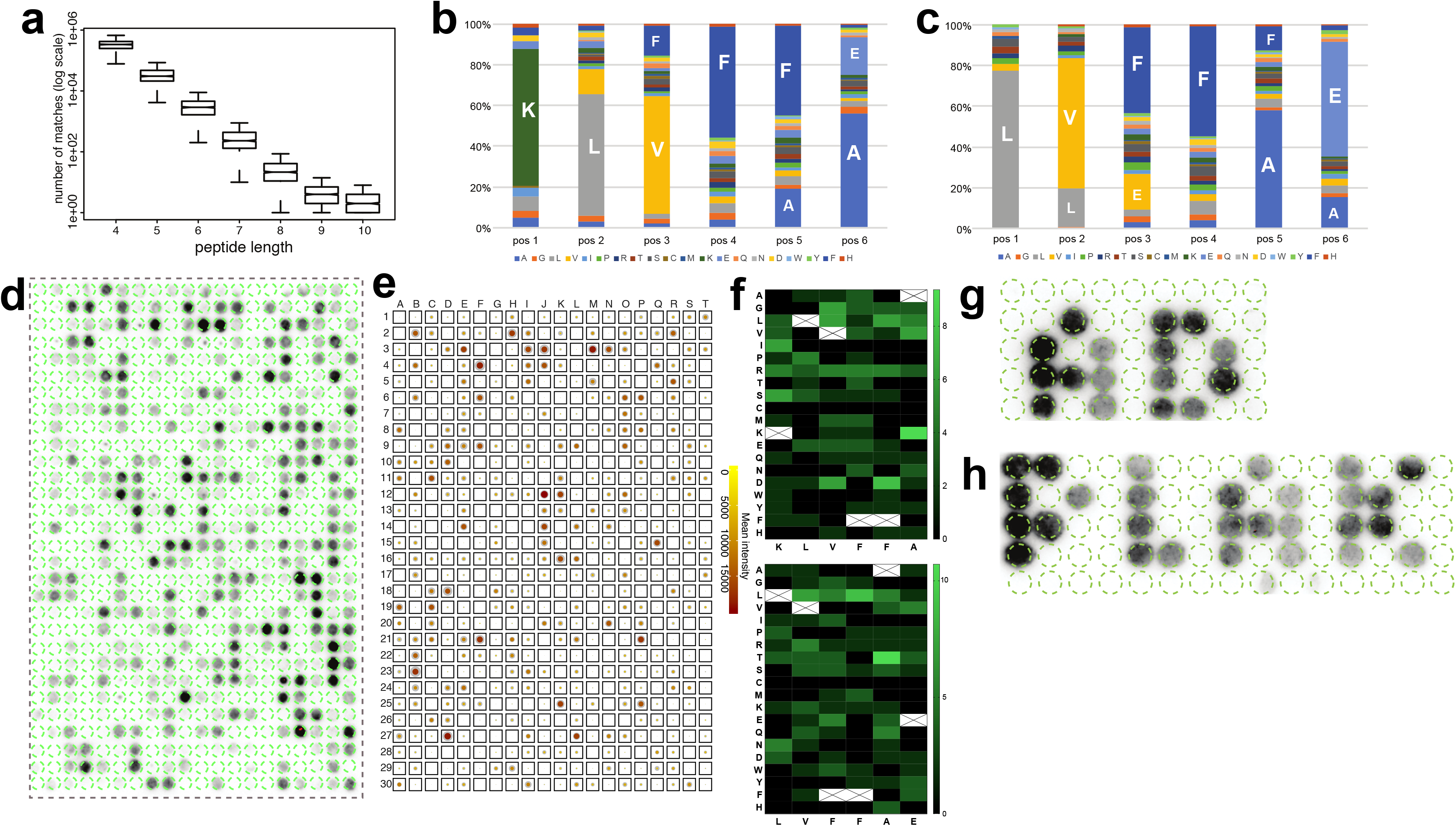
Aβ binding to APR homologues derived from human proteins. a) Sequence similarity in combination to peptide length for 1000 random proteins derived from human proteome. b-c). Distribution of amino acids in homologues to Aβ KLVFFA (b) and LVFFAE (c) proteins. d) Binding of Biot-Aβ1-42 to homologue peptides derived from ~520 randomly selected proteins. e) Summary of Biot-Aβ1-42 binding throughout 8 membranes, color indicates the mean between membranes and the size of the outline the standard deviation. f) Heatmap of amino acid substitutions in membrane hits. g-h) Membrane top binders spell AD and PLAK, white space consists of random sequences.

Since for technical reasons the maximum number of sequences we can currently include on our cellulose array format is 600, we randomly selected this number of fragments from the initial list (Appendix Table S3). We then generated a new membrane with these fragments across the proteome, and to take the immediate sequence context into account, we included 2 N-amino acid and 3 C-amino acid flanks from the matching protein. We exposed this membrane to oligomeric Biot-Aβ1-42 coming from the void fraction of SEC (Figure 1d), and detected the binding pattern to the large membrane using streptavidin-HRP (Figure 2d). This revealed strong binding with some sequences, whereas other showed no or little binding. The binding pattern was reproducible between independently generated replicates of the same membrane and we also generated replicates with the same sequences but in randomized order (Appendix Figure S2). We calculated the overall binders by identifying manually the lowest positive value and calculating its Zscore (named Zcutoff). Every spot with Zscore> Zcutoff in all 8 membranes (at least 2 repeats for 3 randomizations) identified as Biot-Aβ1-42 interactor. This analysis identified 126 consistent binders from this analysis (21%) that bound consistently to oligomeric Biot-Aβ1-42 in all 8 membranes (Appendix Table S4). A summary of the membrane interactions is shown by averaging the binding intensity and standard deviation from 8 membranes (Figure 2e, Appendix Table S4). When we analysed the sequence composition of the bound sequences (Figure 2f), we found that some substitutions were better tolerated than others, eg R in KLVFFA homologues or L in LVFFAE. However, the homologue peptides have single or double mutations to Aβ APRs, which adds a level of complexity in identifying the most favorable mutations.

These experiments show that the presence of specific peptides on the cellulose surface determine where on the membrane oligomeric Biot-Aβ1–42 deposits. To explore this point further, we printed a new membrane in which we spelled the pseudo-words ‘PLAK’ and ‘AD’ using peptide spots from the top 50 binders identified in the previous membranes and surrounded them with peptide spots from random sequences from the human proteome that did not share similarity with Aβ sequences (Figure 2g and h, Appendix Table S5). This confirmed the observations above, that the binding patterns are consistent. Although a cellulose membrane is a poor two-dimensional representation of what may be occurring in a complex tissue such as the brain, these consistent binding patterns show that Aβ amyloids can engage in heterotypic interactions with homologous fragments of otherwise unrelated proteins.

### Heterotypic APR interactions modify Aβ–42 amyloid formation in solution

In order to investigate if the interactions that we detected on the cellulose membrane could affect amyloid aggregation of Aβ1-42 in solution, we generated soluble versions of 32 peptides selected from the membrane (Table 1). We monitored the aggregation kinetics of recombinant rAβ1-42 by ThT fluorescence in the presence of equal amounts (1:1 molar ratio in monomeric units) of these peptides and compared them to peptide alone (Figure 3a,b,c & Appendix Figure S3,S4). We quantified these curves by curve fitting in terms of the lag phase of aggregation (T_lag_), the time at which half the aggregation amplitude is reached (T_1/2_), the total aggregation amplitude (Amp) and the elongation rate (k_e_) (Appendix Table S6). We found that most effects occurred in the lag phase, i.e. where mostly oligomers are populated. We found that 6 peptides showed a statistically significant increase in the lag phase, so slowed down the aggregation of rAβ–42 (Figure 3d), whereas 12 peptides decreased the lag phase, i.e. accelerate rAβ–42 kinetics. In addition, 7 peptides showed significant differences in fluorescence amplitude (Figure 3e).

**Table 1:**
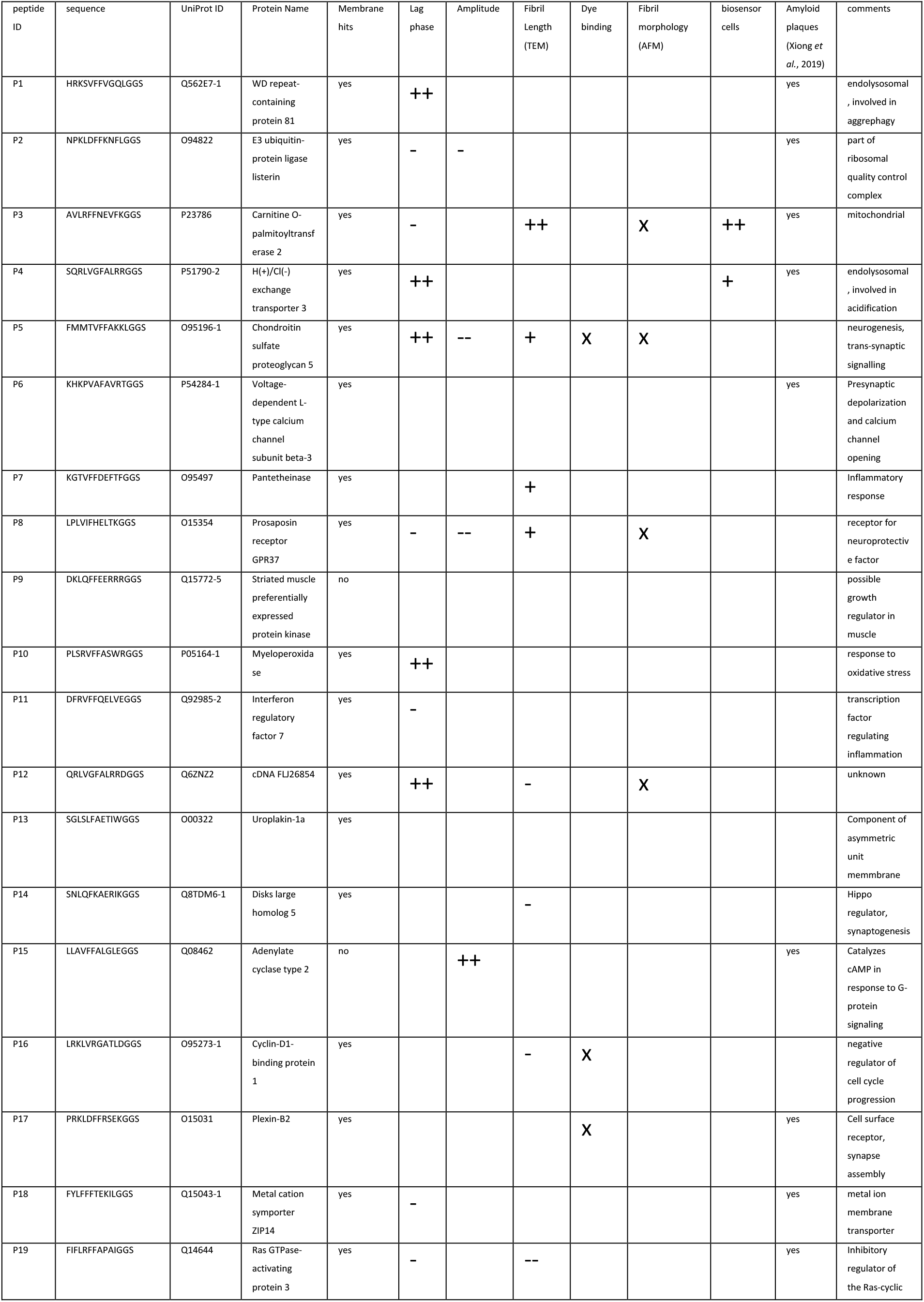
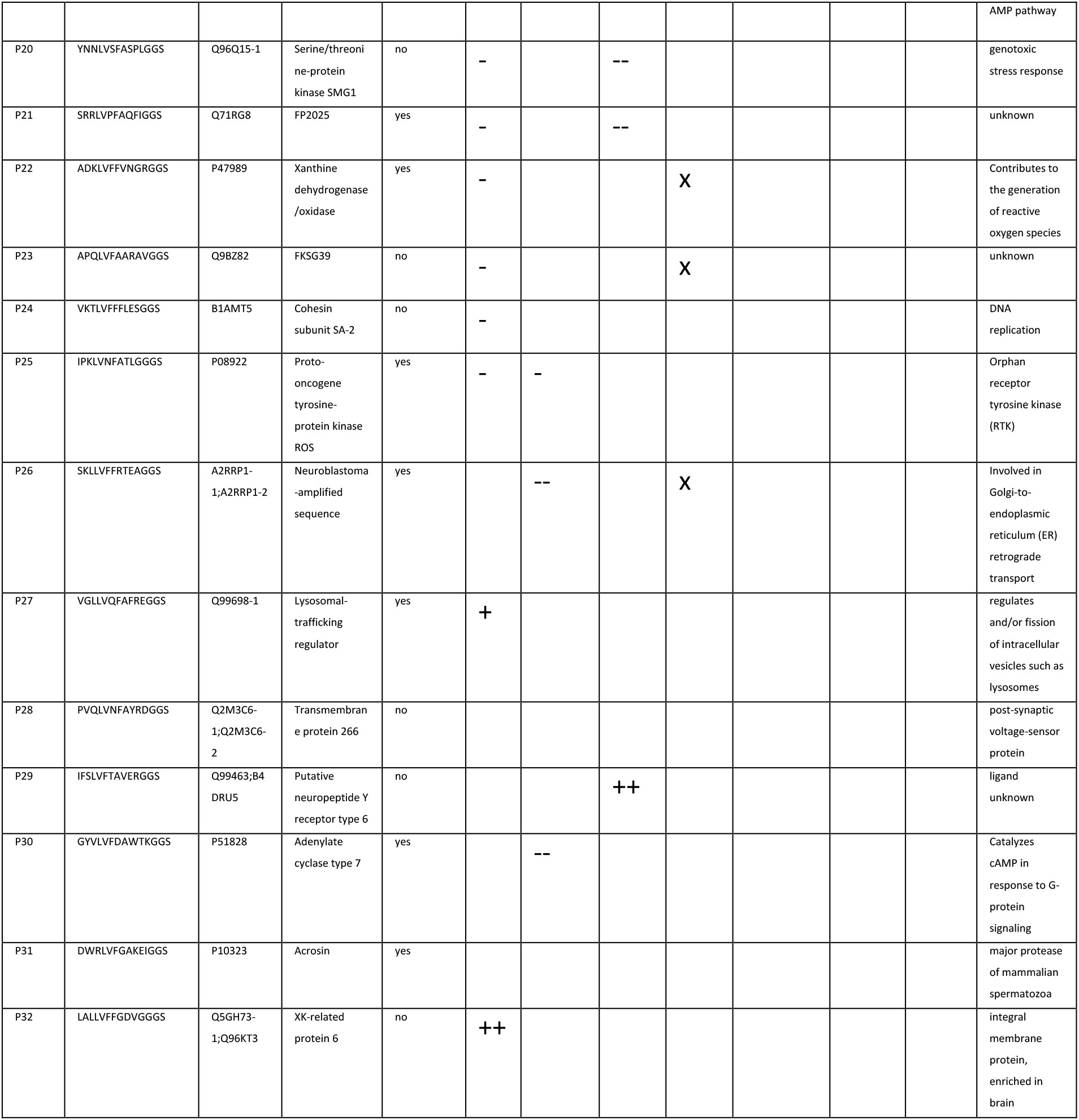
Summary of peptide screen

**3.**
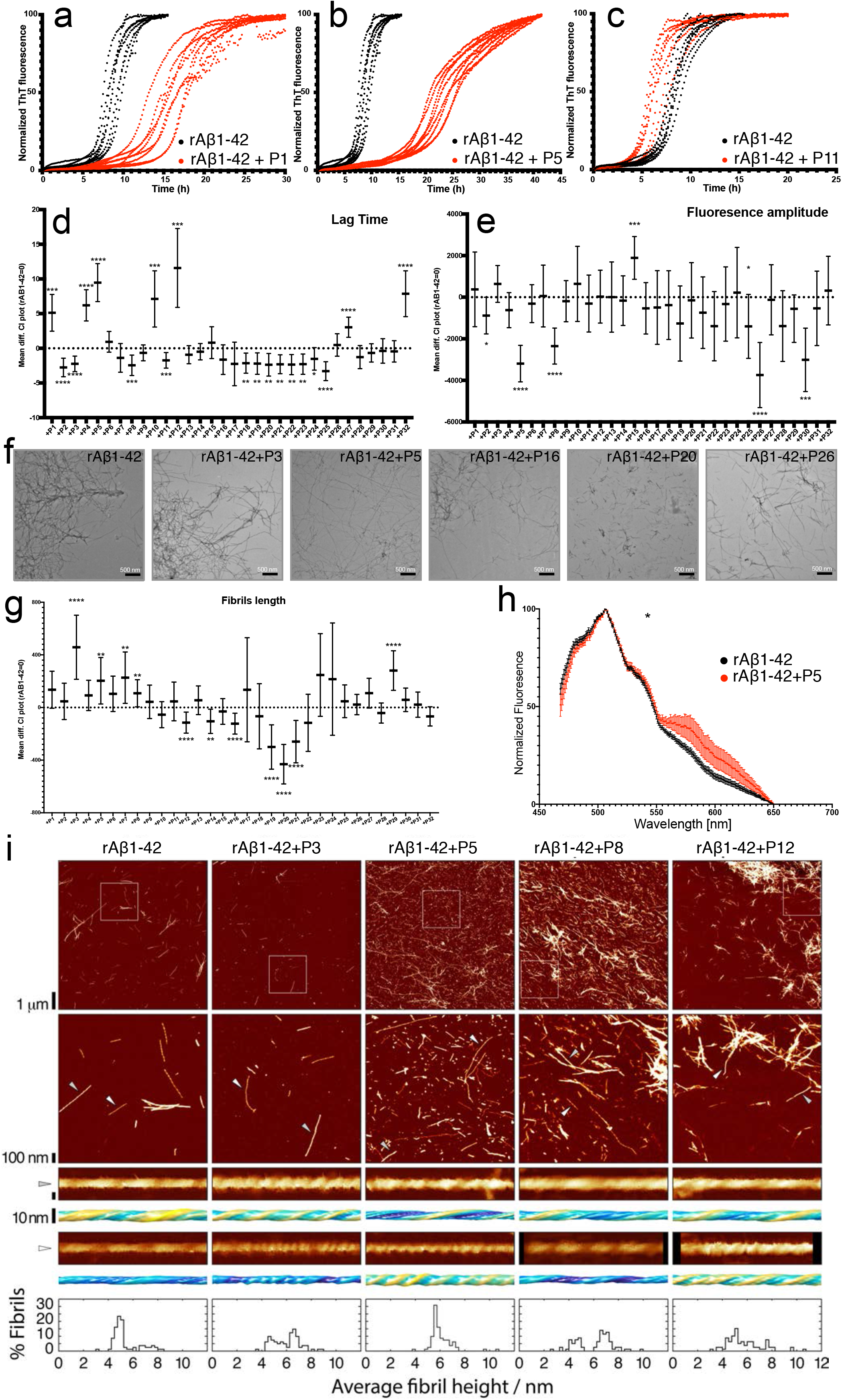
a-c) ThT kinetics of 10uM rAβ1-42 alone or in presence (1:1) of 3 homologue peptides (full screen on Appendix Figure S3,S4, n= 2 independent experiments with 4 repeats) d) Lag time difference between rAβ1-42 alone (rAβ1-42 = 0) and in the presence of peptides (Statistics: Brown-Forsythe and Welch Anova tests with Dunnett T3 multiple comparison corrections) e) Fluorescence amplitude difference between rAβ1-42 alone (rAβ1-42 = 0) and in the presence of the peptides (Statistics: Brown-Forsythe and Welch Anova tests with Dunnett T3 multiple comparisons correction). f) Representative TEM images of fibrils made in presence of 1:1 rAβ1-42:peptides. g) Fibril length difference between Aβ alone and in presence of peptides (Statistics: Brown-Forsythe and Welch Anova test with Games-Howell multiple comparison correction). At least 9 different positions on grid and at least 100 fibrils were counted for each condition (except Aβ+P23, Aβ+P24). Fibril length distribution in Appendix Figure S7. h) Curcumin binding to Aβ fibrils alone or in presence of P5. (n=2, at least 4 repeats, statistics: Kolmogorov-Smirnov test). i) Representative AFM height images of Aβ fibrils alone or in a 1:1 mixture with P3, P5, P8 and P12 peptides are show in the top row. The boxes indicate the magnified regions shown in the second row. Arrows indicate the locations of representative individual fibrils shown in magnified detail, each shown as a 200 nm digitally straightened segment and a 100 nm segment of the corresponding 3D surface envelope model that was calculated from the image data. The scale bar for each row are shown to the left, with both the 3D model and the straightened image data representing 10 nm. The colour scale of the 3D models from blue to yellow indicate the distance (from low to high) between the fibril surface and fibril centre axis to demonstrate their twist patterns. The average fibril height distribution of around 80 manually selected filaments per sample that showed twist patterns characteristic of single, not fragmented, amyloid fibrils are shown in the bottom row.

To investigate the effect of the peptides on the rAβ1-42 mature fibrils, we first analysed the morphology of amyloid fibrils formed in presence of peptides using Transmission Electron Microscopy (TEM, Figure 2f, Appendix Figure S5, S6) and compared it to Aβ fibrils in absence of peptides. We analyzed 10 positions and measured the length of at least 100 fully traced fibrils for each grid to ensure as possible objective quantification. Based on the objective quantification of the fibril length in these images, we found 11 peptides that modified the length of the fibrils (Figure 2g, Appendix Figure S7). Further, we studied the binding to conformationally sensitive amyloid reporter dyes, which alter their emission spectrum depending on the structural detail in the fibril (pFTAA and curcumin, Figures 2h & Appendix Figure S8,S9). Dye binding showed significant differences between rAβ1-42 alone and in presence of peptides for 4 peptides by pFTAA and 2 by curcumin, suggesting a change in the amyloid structure formed (Appendix Figure S9,S10). Moreover, to confirm our observations, we performed Atomic Force Microscopy (AFM) of rAβ1-42 fibrils with a selected number of peptides, which allows the width and morphology of individual filaments to be precisely measured. Interestingly, four of the peptides induced alterations in the mesoscopic arrangement of the rAβ1-42 aggregates as well as the morphologies of individual fibrils (Figure 3i). The presence of the peptides resulted in a change in the width distribution of the fibrils compared to rAβ1-42 alone. The average width increased in the presence of these peptides, which could suggest that some fibrils are composed of greater number of protofilaments or changed protofilament arrangements compared to Aβ alone. Individual fibril surface envelope reconstructions (Aubrey *et al*, 2020; Lutter *et al*, 2020) of well-separated fibrils observed in the AFM images confirm that the morphological details of individual fibril structures formed in the presence of the peptides are indeed different to the rAβ1-42 fibrils formed in the absence of the peptides (Figure 3i), including some fibrils with a higher twist periodic frequency than in the Aβ only sample.

Our data show that peptide fragments of human proteins with local homology to one of the APRs of rAβ1-42 can modify Aβ aggregation kinetics as well as the fibril morphology, under conditions where the two molecules have ample opportunity to interact. It of course remains to be seen whether these peptides still modify Aβ aggregation kinetics in the context of their full-length proteins.

### Proteins containing local homology to Aβ APRs that favour the initiation of Aβ1-42 aggregation in a biosensor cell line

To test the potential effect by full-length proteins on Aβ1-42 aggregation of these heterotypic interactions in complex biological environments, we implemented a simplified model system that allows to investigate the potential of full-length proteins to modulate Aβ aggregation. To that purpose we created a biosensor cell line in HEK293T, in a similar fashion to a previous lines by the Prusiner (Aoyagi *et al*, 2019) and Diamond labs (Kaufman *et al*, 2016), that stably expresses a fusion construct between Aβ1-42 and mCherry tag at N-terminal (Figure 4a). In untreated cells of this line, diffuse mcherry fluorescence is observed throughout the cytoplasm of >95% of the cells. However, when we prepared seeds of rAβ1-42 by sonicating mature amyloid fibrils and added them to the cells by transfection, we observed the appearance of a punctate pattern of the RFP fluorescence (Figure 4b). Automated high content image analysis revealed that the diffuse to punctate transition occurred in a dose-responsive manner (Figure 4c). We also confirmed that the puncta were protein aggregates using Fluorescence Recovery After Photobleaching (FRAP) in a region of increased fluorescence: bleaching of this region resulted in limited recovery supporting that the observed spots were indeed Aβ1-42 aggregates (Figure 4c, Appendix Figure S10).

**4.**
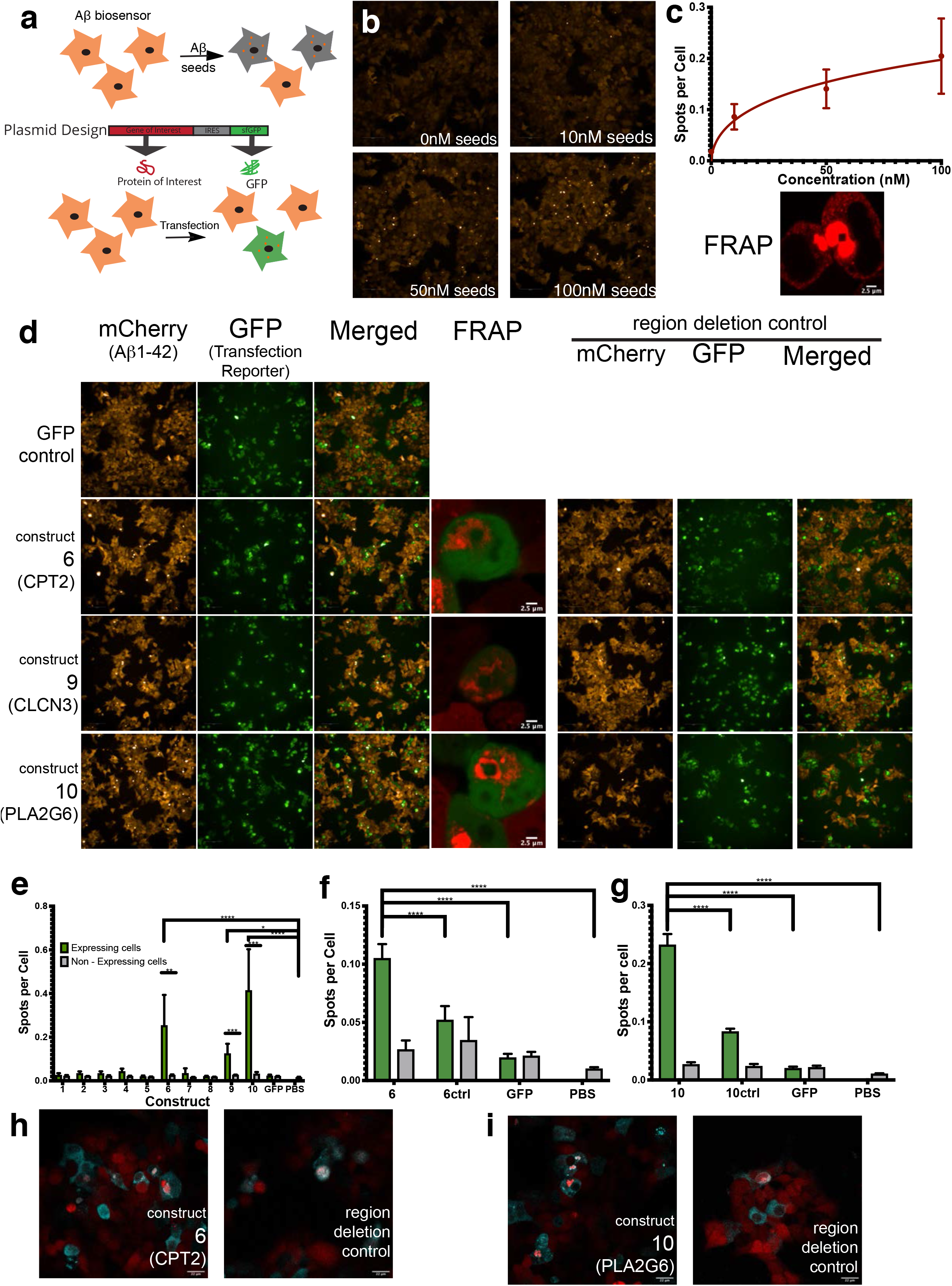
Proteins with homologues to Aβ regions can induce the aggregation of Aβ1-42 in HEK293T cells. a) Experimental setup of inducing aggregation in Aβ biosensor cell line. b) Treating Aβ biosensor with different concentration of rAβ1-42 seeds induces the aggregation of mCherry-Aβ1-42 in a dose dependent matter (n=3 independent experiments, graph: mean and 95%CI). FRAP of Aβ spots shows limited recovery confirming that are indeed aggregates (in detail at Appendix figure S10b,c). d) Representative images of 3 proteins that can induce aggregation of Aβ1-42 in biosensor cell. Increase aggregation is observed in cells expressing the construct but not GFP alone (left panels). FRAP of the resulting aggregates shows no recovery (in detail at Appendix Figure S11). Removal of the homologue regions resulting in reduced aggregation (right panels). (n=3 independent experiments). e) Quantification of number of spots per cell in cells expressing/not expressing the construct (identified by GFP, transfection reporter) (n=3 independent experiments, statistics: ordinary one-way Anova with Dunnett T3 multiple comparison correction, unpaired t-test for transfected/non-transfected cells). Bar plot: mean with 95%CI. f-g) Quantification of Spots per cell for construct6 and construct6 control (6Ctrl, removal of homologue region) (f) and construct10 and construct10 control (10ctrl, removal of homologue region) (g). (n=4 independent experiments, statistics: ordinary one-way Anova). Bar plot: mean with 95%CI. h-i) Confocal images of colocalization of protein of interest with Aβ aggregates. (Contrast of images were enhanced to 0.1% saturated pixels using Fiji).

In order to test if proteins with homologous regions to the Aβ APR segments are capable of inducing Aβ1-42 aggregation in a similar way that seeds did, 10 expression constructs were generate containing each gene of interest as well as a fluorescent reporter (GFP) expressed by an Internal Ribosome Entry Site (IRES) (Figure 4a). This setup allows us to quantify Aβ1-42 aggregates in cells that are expressing our protein of interest, and compare it directly with non-transfected cells from the same well. The 10 proteins were chosen based on their binding signal in our peptide microarrays, their synthesis potential, and/or their connection to brain or neurodegenerative diseases (Table 2). From those 10 constructs, 8 correspond to the full-length protein and 2 had to be cut slightly due to size limitations of the synthesis method (TWIST bioscience). When we transfected the biosensor cells with these constructs and compared the number of spots per cell in transfected and non-transfected cells, we identified 3 proteins (correspondent to constructs 6, 9 and 10) that by simply being overexpressed induced a significant increase of Aβ1-42 aggregation (Figure 4d&e). As further controls we ensured that we did not observe induction of puncta in mock transfected cells (PBS), nor in cells transfected with a control plasmid expressing only GFP. Moreover, FRAP showed that also these spots were Aβ1-42 aggregates since no recovery was observed after bleaching (Figure 4d, Appendix Figure S11).

**Table 2:**
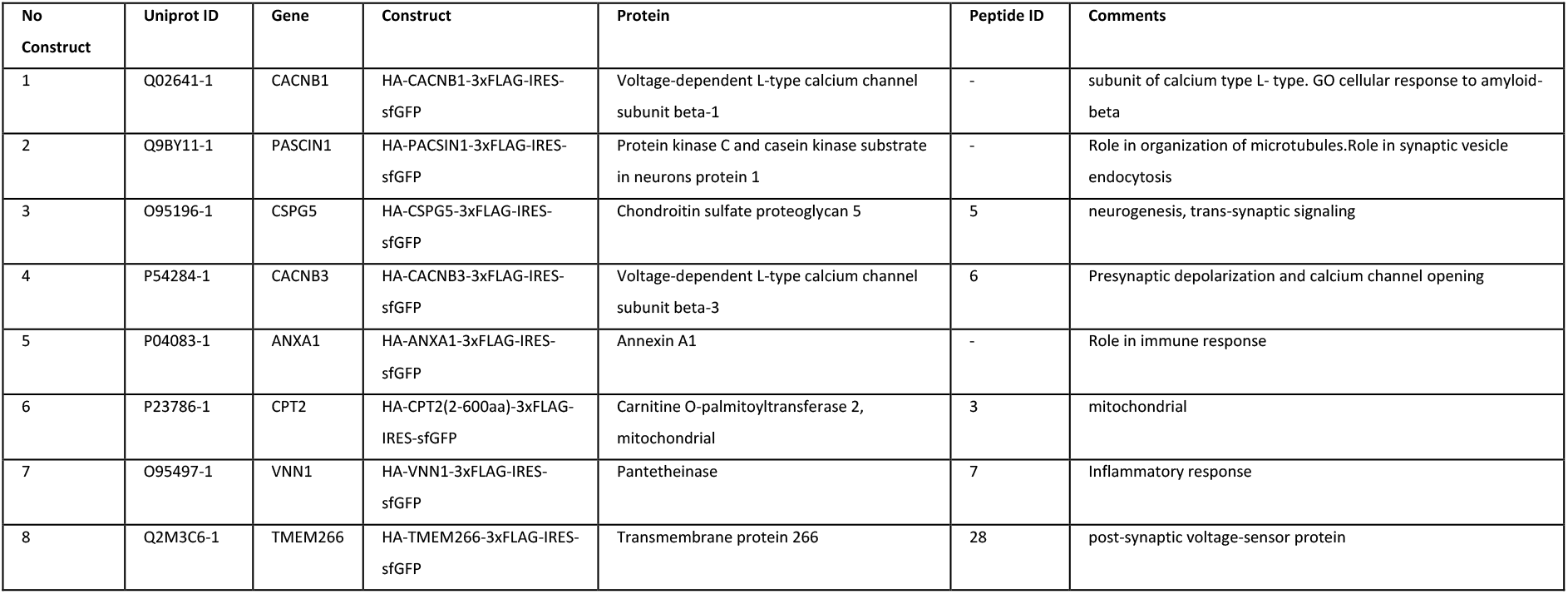
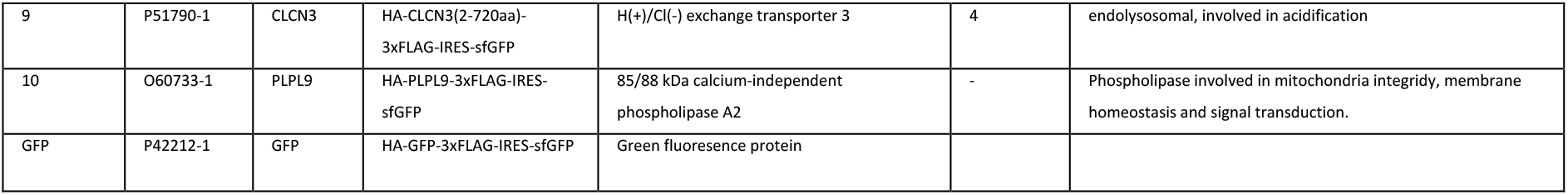
Summary of constructs used in biosensor

To test our hypothesis that the effects we observed on Aβ1-42 aggregation were due to the presence of the sequence segments that are homologous to Aβ APRs, we synthesized additional constructs in which we deleted 10-60 aa containing the Aβ APR homologues. We tried very short deletions, corresponding to the homologous segment, but we also made larger deletions. The latter is because APRs are typically part of the hydrophobic core of a globular folded domain, and hence there are typically 1-3 other elements of the structure that have been evolutionarily optimized to interact with the APR. Hence, deletion of just the APR promotes 3D domain swapping type of interactions, where the APR in Aβ would interact with the remaining compatible regions in the rest of the domain, which we have previously shown promotes aggregation through that mechanism (Rousseau *et al*, 2001). Construct 6 expresses 1-600aa of CPT2 protein, containing the peptide P3 from Table 1 that significantly affected the kinetics and morphology of Aβ1-42 aggregation (Figure 3) and as a full-length protein increased the aggregation of Aβ1-42 in our biosensor cell line. We design a control construct, 6Ctrl by removing the homologous region (aa 379-388) and expressed in Aβ1-42 biosensor. The absent of homology significantly decreased Aβ aggregation when compared to initial construct (Figure 4d,f). Moreover, immunofluorescence confirmed that the majority of aggregates exist in cells expressing CPT2 and partially colocalized with Aβ aggregates (Figure 4h). However, that was not the case for the control (Figure 4h). PLA2G6 (Construct 10) showed the most acute increase of Aβ1-42 aggregation in our biosensor. PLA2G6 control, 10Ctrl, made by removing ANK7 domain (aa 349-378). Expression of this control in Aβ biosensor reduced significantly the aggregation of Aβ1-42(Figure 4d,g). Indeed, presence of PLA2G6 induced the aggregation of Aβ1-42 as seen by immunofluorescence. Moreover, PLA2G6 increased signal is observed in Aβ1-42 aggregates (Figure4i). This is not observed in control (Figure 4i). Finally, the third protein that induced aggregation of Aβ1-42 is CLCN3 (Construct 9) (Figure4d,e). Two controls, Ctrl1 and Ctrl2, was made by removing a 63aa domain and 20aa respectively. Removal of the homologous region only showed a minor decrease in the aggregation of Aβ1-42 (Appendix Figure S12a,b). Moreover, we could not detect CLCN3 in the Aβ1-42 aggregates by immunofluorescent (Appendix Figure S12c), and we found another homologous region elsewhere in the CLCN3 sequence, further complicating the analysis. This suggests that the effect of CLCN3 expression on Aβ1-42 aggregation that we observed is indirect or only partially resulted from heterotypic amyloid interactions.

### Proteins with local sequence homology to Aβ APRs are enriched in human Aβ plaques

In order to study if proteins containing segments with high sequence similarity to the APR regions of Aβ42 may be enriched in Alzheimer’s disease-associated amyloid plaques, we analysed a proteomics dataset of hippocampal amyloid plaques of AD patients generated from Xiong et al (Xiong *et al.*, 2019). This study provides high quality proteomics profiling at unsurpassed depth of amyloid plaques (AP) and, importantly, nearby control tissue, obtained by tag-labelling high-throughput mass spectrometry, which is a highly quantitative method. We searched the proteins identified in amyloid plaques (1125 AP proteins) and adjacent non-plaque regions for segments with sequence homology to Aβ42 in an unbiased manner: We divided the Aβ sequence into hexapeptides using a sliding window approach and searched the proteins for homologous segments, allowing up to two mutations. To identify if some Aβ segments were overrepresented in amyloid plaques, we studied the occurrence of homologous segments to each Aβ region in amyloid plaques and compared to the control region proteins to calculate enrichment values. Our analysis found six hexapeptide segments of Aβ to be overrepresented in AP proteins, compared to tissue proteins (Figure 5a). Interestingly, those positions are nearly perfectly overlapping with the APR regions of Aβ APR, as would be expected if the enrichment had resulted from heterotypic APR-interactions. Two of these hexapeptides reside in the central region, and partially cover the KLVFFA APR, and 4 additional hexapeptides reside in the C-terminal APR. To further test if the observed overrepresentation is caused by biases in the background proteins, or form set-size imbalances between the AP and control sets, we employed random sub-sampling to estimate the distribution of homologous regions in the background (in a so-called bootstrapping approach). We used the proteins identified in the tissue and created 1000 random samples with protein numbers equal to the AP proteins (Figure 5b). In a similar way, we also created 1000 random samples of the same size taken from the whole human proteome and a random plaque sample for each sample (Figure 5c). Both controls showed that the regions identified to be overrepresented in the plaque are residing in the tails or well outside of the random distributions, supporting the notion that the enrichment did not occur by chance. In fact, the bootstrapping approach suggests there may be more over-represented regions, but we took a conservative approach and only considered regions that withstood all statistical testing.

**5.**
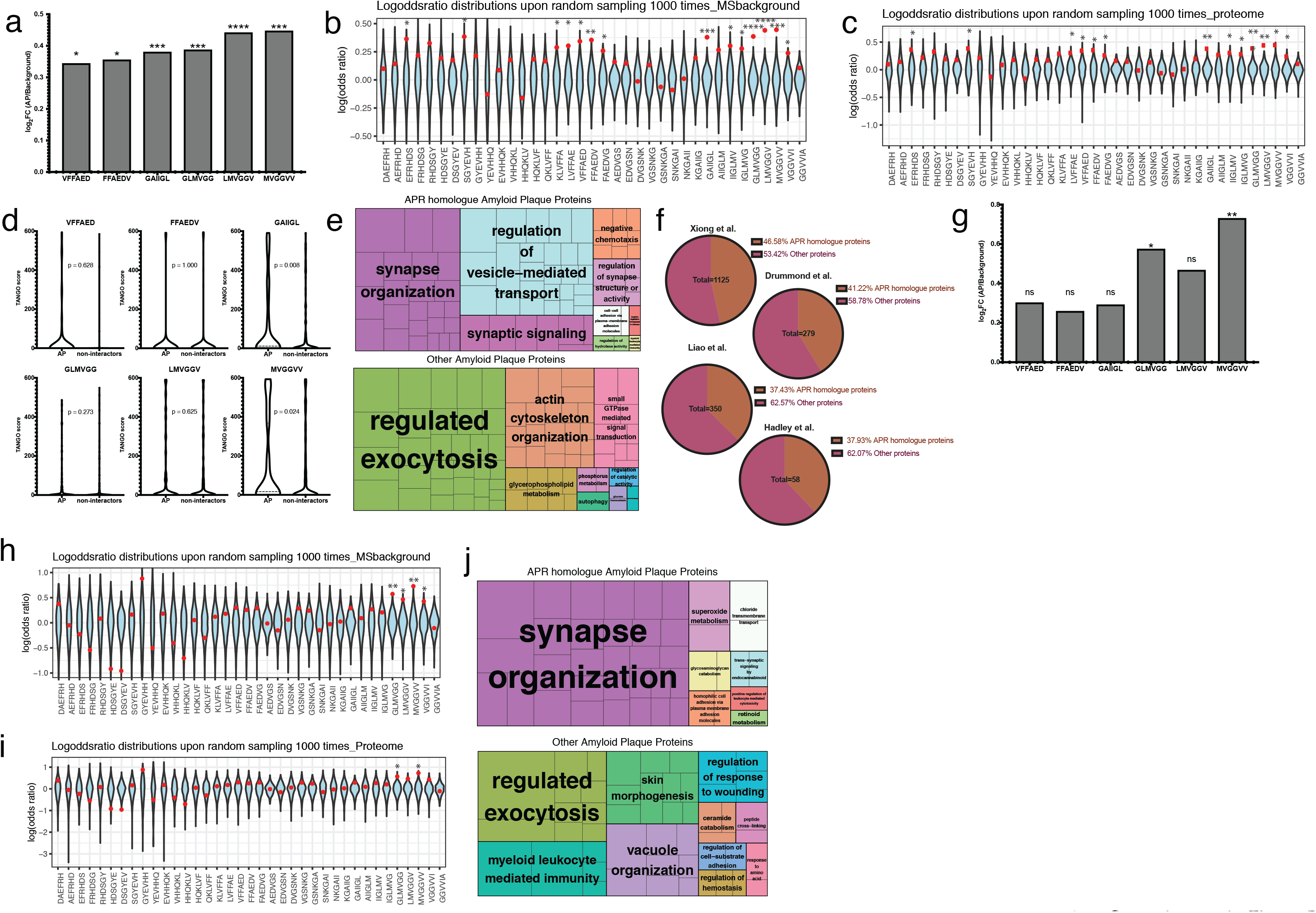
a) Overrepresentation of Aβ APRs in amyloid plaques of AD brains (hypergeometric test with Bonferroni correction) b) Log-odd ratio of random sampling from mass spectrometry background. Red dot indicated the true values of analysis c) Log-odd ratio of random sampling from human proteome. Red dot indicated the true values of analysis d) TANGO scores of homologue APRs in amyloid plaques and non-amyloid plaques proteins (statistics: Kolmogorov-Smirnov test) e) Biological pathways enrichment of Aβ APR homologue related and non-related proteins derived from AD amyloid plaques f) APR homologue proteins identified in other MS studies g) Overrepresentation of Aβ APRs in amyloid plaques of nonAD brains. h) Log-odd ratio of random sampling from mass spectrometry background. Red dot indicated the true values of analysis i) Log-odd ratio of random sampling from human proteome. Red dot indicated the true values of analysis j) Biological pathways enrichment in nonAD amyloid plaques.

After we identify the regions that are overrepresented in amyloid plaques, we wondered if the AP proteins containing homologous segments to the Aβ APRs had a higher aggregation propensity than proteins found in the control tissue that also contain homologous segments, but that are not found in the plaques. To do so we used the TANGO algorithm to analyse the protein segments with homology to Aβ regions identified before as overrepresented. Our analysis showed that homologous regions from AP proteins showed a higher aggregation propensity than the ones not found in the plaques, with two regions (GAIIGL and MVGGVV) showing statistically significant differences (Figure 5d). These results suggest that again that heterotypic APR interactions may be involved in the enrichment of these proteins in the plaques.

Because proteins associated with amyloid plaques may be involved in high risk pathways for AD, we sought to identify the pathways that proteins with homology to Aβ APRs are involved. The two previous groups of amyloid plaques were searched against Gene Ontology Biological process pathways and the significantly enriched pathways were isolated (Figure 5e). Interestingly, AP proteins with homology to Aβ APRs were found to play role in “synaptic organization, structure and activity”, pathways highly relevant to AD, since synaptic dysfunction is known to play an important role in AD progression. Finally, we wanted to test if a similar occurrence of homologue to Aβ APR proteins exists in other proteomic studies of AP. Indeed, a similar trend is observed in other AP proteomic profiles, with a range of 35-45% of proteins found in APs to have a sequence homology to Aβ APRs (Figure 5f)

To investigate if a similar overrepresentation is seen in amyloid plaques from non-AD brains and APP/PS1 mouse model, we analysed them in a similar way (Figure 5g, 6a). In the case of amyloid plaques from non-AD brains, two of the 6 previously identified regions were found to be overrepresented (Figure 5g,h,i). Since two of the previously identified six positions are found to be overrepresented, we hypothesized that proteins may still be interacting through the other 4 positions but without reaching the high levels observed in AD brains yet. So, we used the AP proteins found in non-AD plaques that have homology to all 6 regions and the other AP proteins to do the enrichment analysis of Gene Ontology biological processes, like previously. Interestingly, AP homologue proteins of non-AD brains were also found to be involved in synaptic pathways (Figure 5j).

Furthermore, we analysed proteomic data of amyloid plaques from APP/PS1 mouse AD model obtained using a similar method as the human amyloid plaques(Xiong *et al.*, 2019). To do so we analysed in a similar way as previously two biological replicates. Remarkably, the overrepresentation of Aβ homologue regions was completely abolished in both replicates (Figure 6a). However, the random distribution identified 2 to 3 positions slightly significantly overrepresented but not in the extend observed in human AD brains (Figure 6b,c). Mouse models usually are overexpressing Aβ which leads to rapid aggregation and deposition compared to the slow process found in humans. So, the lack of overrepresentation in mouse model suggests that self-aggregation in promoted in the mouse model, reducing the opportunity for heterotypic interactions, thereby potentially explaining the differential toxicity that is observed between humans and mouse.

**6.**
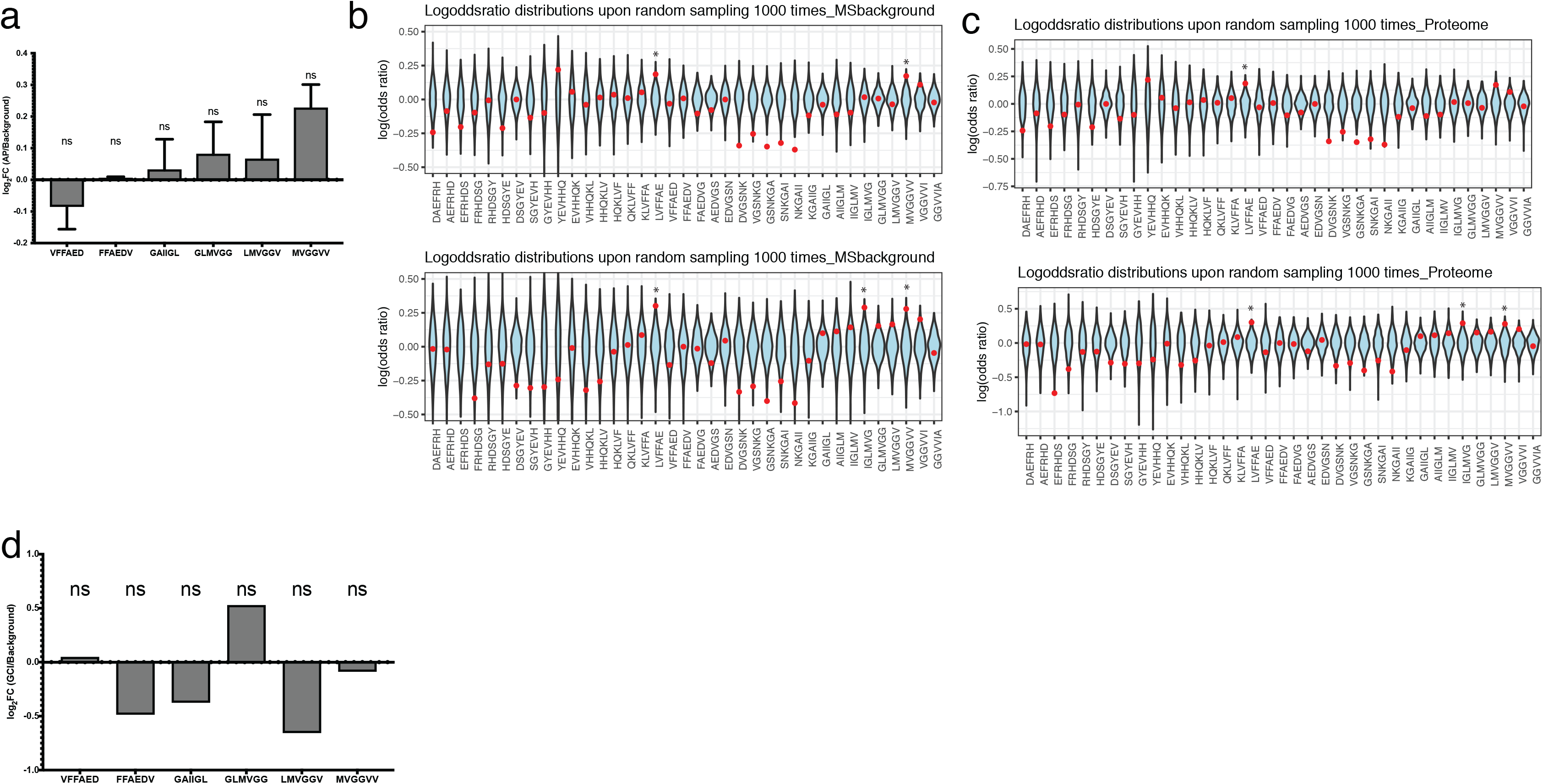
a) Overrepresentation of Aβ APRs in amyloid plaques of APP/PS1 mouse brains (mean ±SD) from two biological replicates. b) Log-odd ratio of random sampling from mass spectrometry background for both replicates. Red dot indicated the true values of analysis c) Log-odd ratio of random sampling from mouse proteome for both replicates. Red dot indicated the true values of analysis. d)No overrepresentation of Aβ APRs was observed in proteins from Glial cytoplasmic inclusions.

Finally, to test if the observed overrepresentation is exclusively seen in amyloid plaques and not aggregates from other proteins, we sought to analyse proteomic data from other pathological aggregates. We chose to analyse a study of Glial cytoplasmic inclusions (GCIs), which are mainly composed from α-synuclein, from Multiple system atrophy (MSA) brains (McCormack *et al*, 2019). These GCIs were isolated from Basal Ganglia of 5 MSA brains and the proteins identified in at least 4 cases were used as the aggregation-related proteins. Since, this analysis comes from purification of aggregates, no normal tissue was analysed. To overcome this problem, we used as tissue, proteins identified in a proteomic study of Basal ganglia (Fernandez-Irigoyen *et al*, 2014). From our analysis, no overrepresentation of any Aβ region was observed in those α-synuclein enriched aggregates. This result indicates that the proteins with Aβ homology regions are primarily found in amyloid plaques and are not significantly overrepresented in aggregates driven by other proteins.

## Discussion

Understanding selective neuronal and regional vulnerability requires knowledge of both loss- and gain-of-function effects associated to amyloid deposition. Much of our understanding on the role of amyloids in neurodegenerative diseases derives from improvements in our knowledge of the native function of these proteins. At the same time, it remains hard to contextualize the role of amyloid deposition and in particular the specific interactions they engage and how these contribute to disease. Historically, a lot of the mechanistic thinking on the role of amyloids in disease was inspired by their common structural properties and the presumption that amyloids therefore also possess generic modes of interaction with their environment (Bucciantini *et al*, 2004; Bucciantini *et al*, 2002; Campioni *et al*, 2010; Flagmeier *et al*, 2020). Yet the overall view that emerged is that of rather promiscuous amyloids that easily interact with various lipids & membranes or nucleic acids and that co-precipitate in unspecific manner with many other proteins (Olzscha *et al*, 2011). This view has been complemented by the realization that protein expression of many proteins – and particularly in the brain – leads to their supersaturation (Ciryam *et al*, 2015; Tartaglia *et al*, 2007) while proteostatic regulation erodes with ageing (Labbadia & Morimoto, 2015) together setting the scene of a metastable proteome that becomes increasingly prone to collapse. While these finding have vastly increased our understanding of the existence of misfolding & aggregation diseases and the conditions favouring their development they do not explain the specific neuronal vulnerabilities characterizing each of these diseases.

We here show that Aβ oligomers can interact with various short APR homologous segments of otherwise unrelated human proteins and that such interactions modify Aβ aggregation kinetics and fibril morphology. Such heterotypic interactions *de facto* modify the cellular vulnerability of a reporter cell line for spontaneous Aβ aggregation. While this cellular model does not aim to mimic the pathological context of amyloid initiation in Alzheimer’s disease it does in a simplified manner illustrate how cellular vulnerability for amyloid initiation can be shaped by specific interactions of a disease amyloid with its proteomic background. While the cross-β propensity of APRs favours self-assembly it also allows some degree of ‘off target’ interaction which can further facilitate or inhibit amyloid assembly thereby sensitizing or protecting cells from amyloid nucleation. Our findings are supported by the increasing observation of heterotypic amyloid assembly both in disease and for the functional regulation of biological processes (Louros *et al*, 2016; Zhou *et al*, 2012). These experiments have further revealed the importance of local sequence homology in heterotypic amyloid assembly, indicating that the selectivity of these interactions is related to the sequence similarity level shared between constituent elements (Kehrloesser *et al*, 2016; O’Nuallain *et al.*, 2004; van der Kant *et al.*, 2021; Wang & Fersht, 2015; Xu *et al*, 2011; Yan *et al*, 2007). Co-assembly often results in accelerated and more severe pathological outcomes, as reported for Aβ and α-synuclein in AD and PD patients (Mandal *et al*, 2006). Both proteins have been shown to hold a nucleation effect towards Tau (Colom-Cadena *et al.*, 2013), whereas huntingtin may as well be involved in cross-fibrillation mechanisms, leading to polyglutamine disorders (Furukawa *et al*, 2009). Heterotypic assembly has also been associated to amyloid transmissibility of neurodegenerative disorders and as a causative agent for the progression of certain forms of systemic amyloidosis (Westermark & Westermark, 2010).

The observation that heterotypic interactions affect the mesoscopic structure of Aβ fibrils in vitro suggests that such interactions can potentially also contribute to polymorphic bias in disease. Thus, next to specific posttranslational modifications and interaction with non-proteineous prosthetic ligands observed in amyloid cryoEM structures, heterotypic amyloid interactions could represent yet another way in which amyloid polymorphism can be affected by and possibly also affect the specific environment in which they are formed (Arakhamia *et al*, 2020; Scheres *et al*, 2020; Wesseling *et al*, 2020).

To evaluate whether heterotypic interactions occur in a neurodegenerative context we re-evaluated deep proteomics data of human Aβ plaques (Xiong *et al.*, 2019). We found that sequences homologous to Aβ APRs are enriched in plaques while this is not the case for non-APR segments of Aβ. While Aβ plaques are of course mostly composed of Aβ itself, the enrichment of these ‘contaminants’ suggests that heterotypic interactions occur in the process leading to plaque formation. Interestingly we do not find such enrichment in the Aβ overexpression APP/PS1 mouse model. Possibly this could mean that in this case overexpression is the dominant driver of plaque formation thereby outcompeting any possibility for heterotypic interaction. When analysing the function of heterotypic plaque components, we find that they mainly cluster in gene ontologies related to synaptic regulation and the regulation of vesicle-mediated transport suggesting that heterotypic interactions observed in plaques are associated to the synaptopathology of Alzheimer’s disease (Forner *et al*, 2017). This raises the question of the nature of such association. Does heterotypic plaque composition reflect interactions that follow synaptic damage or do such interactions directly participate in the pathological chain of events resulting in synaptic breakdown? In the latter case the gain-of-function effects of heterotypic interactions could be bidirectional whereby protein interaction with Aβ facilitates Aβ initiation while Aβ also perturbs the normal function of these proteins. Previous work with synthetic amyloids (Betti *et al*, 2016; Gallardo *et al*, 2016; Michiels *et al*, 2020), the inhibition of homologues by p53 tumor suppressor (Xu *et al.*, 2011) or the inhibition of mammalian necroptosis by viral proteins supports the possibility that heterotypic amyloid interactions (Pham *et al.*, 2019) can result in the functional knock down of target proteins. It is also probable that heterotypic interactions will be favoured in situations where proteins are (partially) unfolded e.g. during translation, translocation or due to physiological ageing implying additional spacial and temporal context.

It remains to be seen how the net result of the simultaneous expression of many homologous sequences adds up and to what degree these contribute to shaping neurodegenerative diseases. It also remains to be explored whether and how heterotypic amyloid interactions relate to genetic risk factors and to the complex pathophysiologic alterations observed in amyloid-associated neurodegenerative diseases. For now, we here present evidence for a generic molecular mechanism predisposing Aβ amyloid structures to local sequence-specific gain-of-function binding interfaces allowing them to interact with specific proteins in a complex proteome. Thereby, these proteins affect the aggregation of Aβ which in turn may functionally affecting these proteins by promoting co-assembly in amyloid deposits.

## Materials and Methods

### In silico screening of Aβ homologous APR

The human proteome (UniProt, 2008) was computational screen for protein sequence fragments that are highly similar to the APR of the Aβ peptide (KLVFFA and LVFFAE), allowing maximum 2 mismatches within a hexapeptide. An inhouse generated algorithm was used to identify these homologue sequences. Approximately 600 homologous peptides were randomly picked from the human genome and further in-house synthetized/printed in membrane peptide arrays.

### Homologous peptide membrane arrays and Aβ1-42 binding assay

The peptide arrays were developed through SPOT synthesis on acid stable cellulose membranes using the Intavis Multipep RSi synthesis robot. The peptides were synthetized, from C-terminus to N-terminus, starting with a GGS linker and containing a PEG spacer (Aims-Scientific). The obtained peptide array membranes were first incubated in 50 % methanol for 10 min, followed by three short washes in PBS-T (PBS, 0.05% Tween-20). The membranes were blocked overnight in 1% BSA in PBS-T, then washed for three times 5 min first in PBS-T and next in the incubation buffer (10 mM MES, 150 mM NaCl, 0.05% Tween-20, pH 5.5). Then the membranes were incubated with Biot-Aβ1-42 in incubation buffer supplemented with 100 mM threhalose, for 1 h at room temperature.

The used Biot-Aβ1-42 sample was obtained by solubilizing 0.1 mg Biot-Aβ1-42 (rPeptide) in HFIP for 1 h, followed by 10 min water bath sonication, and finally drying under a N_2_ stream. The film was re-dissolved in 8 M urea or 7M GnHCl in 50 mM Tris, pH 7.4 and the sample was run over a Superdex 75 Increase 10/300 GL gel filtration column (GE Healthcare) equilibrated on the same buffer supplemented with or without 150 mM NaCl, 8M Urea respectively. The void peak of Biot-Aβ1-42 was diluted 1:32 in the incubation buffer, with a final concentration of 0.25 M urea. 10μM of monomeric Biotin-Aβ42 was left to aggregate while measuring the ThT kinetics and 100nM of monomeric or different aggregating species were incubated in the membrane.

After the incubation with Biot-Aβ1-42, membranes were washed four times 5 min in Aβ incubation buffer, then in PBS-T, and then incubated with Streptavidin-poly HRP (Pierce 22140), diluted 1:100,000 in PBS-T, for 1 h. Finally, membranes were washed in PBS-T for three times 5 min and developed through chemiluminescence using a ChemiDoc XRS (Bio-Rad).

### SEC-MALS analysis

The MW of the Biot-Aβ1-42 sample used in peptide membrane assays was studied using multi-angle light scattering (MALS) on a DAWN HELEOS MALS instrument from Wyatt Technology (Santa Barbara, CA, U.S.A.) with an incident laser wavelength of 658 nm. The proteins were separated using a Superdex 75 Increase 10/300 GL gel filtration column (GE Healthcare) connected to an LC-10 Prominence HPLC system (Shimadzu), equilibrated with 50 mM Tris, pH 7.4 containing 150 mM NaCl, at a flow of 0.3 ml/min at RT. First, 25 μL of a 2.0 mg/mL bovine serum albumin standard (Pierce) was injected. The scattering intensities at different angles were collected, corrected for the refractive indices of glass and solvent and normalized using the standard. Then, a 0.1 mg Biot-Aβ1-42 HFIP film was re-dissolved in 250 μL 8 M urea in 50 mM Tris, pH 7.4, passed through a 0.2 μm Spartan filter (Whatman) and 100 μL was injected on the column. The value of d*n*/d*c* (wherein *n* is the refractive index of the solution and *c* the solute concentration) was set to 0.185 mL/g and the scattering data (collected at an interval of 0.5 s) were then fitted according to Zimm formulation.

### Membranes analysis

The signal for each spot in membrane was quantified using ImageLab (Biorad). 100 spots in the borders of the membrane was also quantified and the mean and SD of the background was calculated. We identified manually the lowest positive value for each membrane and calculated the 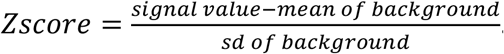. This Zscore was rounded up in an attempt to exclude the faint spots and labeled as Zcutoff. Each Z value of spots was calculated and the ones higher than the Zcutoff identified as hits. Hits consistent in all 8 membranes were identified as interactors of Aβ.

### Purification of Met-Aβ1-42 (rAβ1-42)

The purification of the recombinant Met-Aβ1-42 peptide was performed in-house based on the previously reported protocol (Walsh *et al*, 2009), using the human Met-Aβ1-42 expression plasmid, a kind gift from C. Gomes (FCT, Lisbon). Briefly, the Met-Aβ1-42 plasmid was over-night expressed in *E. coli*, and used to inoculate 1L of M9 culture medium, freshly supplemented with 50 mg/ml ampicillin and chloramphenicol, 2mM MgSo_4_, 0,1mM Ca_2_Cl and 20% glucose. After reach an OD600: 0.6-0.8, the expression of the plasmid was induced with 0.5M IPTG and leave to grow for 4 h. The collected pellet was suspended in 10mM Tris, 1mM EDTA pH8, sonicated and centrifuged at 16000rpm and the final pellet dissolved in the same buffer supplemented with 8M. After diluted to 2M urea, a first ion-exchange chromatography was performed in a DEAE Sepharose resin (GEHealthcare), using the suspension buffer supplemented with 25mM NaCl, as a binding buffer, and with 125mM NaCl as elution buffer. The purified solution was filtered in 30KD spin columns (GEHealthcare), further concentrated in a 3KD ones and the final sample lyophilized in vials with 1mg or 0.65mg.

Prior to each experiment, the lyophilized sample was suspended for 1 h at room temperature in 800ul of 7 M GuHCl in 50mM Tris pH8, centrifuged 5 min at 15,000 rpm at 4°C, and the supernatant injected (using 1 ml injection loop) in a Superdex 75 10/300GL gel filtration column (GE Healthcare), previously equilibrated with 50mM Tris pH8 buffer. The fraction containing monomeric Met-Aβ1-42 was collected and kept on ice, the concentration was determined in a NanoDrop 2000 (Thermo Fisher Scientific), using a molecular weight of 4645 Da and an extinction coefficient of 1.49. The sample was immediately used in several assays.

### Seeds

Biot-Aβ1-42 or rAβ1-42 seeds were obtained by sonicating mature fibrils for 15min (30sec on,30 sec off) at 10°C using Bioruptor Pico.

### ThT kinetic Assay

10uM of monomeric Biot-Aβ1-42 in 50mM Tris pH7.4 was pipetted to μclear medium binding half area plates (Greiner, #675096) and ThT was added to a final concentration of 25 μM. ThT binding was measured over time (through excitation at 440 nm and emission at 480), using a Fluostar fluorescence plate reader (BMG Labtech) at 30°C. ThT kinetics for biotin-Aβ42 was done in a similar way by adding 5 or 10% of biotin seeds.

The monomeric samples of Met-Aβ1-42 obtained after the purification protocol, described above, with a concentration of 10 μM was incubated at 30°C with a constant shacking for 4 days.

All used peptides were purchased from Genscript, and in their design scheme have a GGS on the c-terminus, a PEG2 on both terminus and are acetylated and amidated, respectively on N and C terminus. Peptides were solubilized in HFIP, aliquoted in 0,25mg vials, dried under N_2_ stream and stored at −20°C. Peptide HFIP films were dissolved in 50 mM Tris pH 8 buffer, filtered through a 0.22 um Millex-GV spin filter and diluted to 20 μM. In-house purified Met-Aβ1-42, described above, was diluted to 20 μM. Peptide and Met-Aβ1-42 were mixed 1:1 with a final concentration of 10 μM each, in 50 mM Tris pH8 buffer. Mixtures containing only peptide or Met-Aβ1-42 were used as controls.

The mixtures of Aβ1-42 and the peptides were pipetted to a μclear medium binding half area plates (Greiner, #675096) and ThT was added to a final concentration of 25 μM. ThT binding was measured over time (through excitation at 440 nm and emission at 480), using a Fluostar fluorescence plate reader (BMG Labtech) at 30°C, with a readout every 10 minutes and 10 seconds of shaking before each readout. Similar peptide:Met-Aβ1-42 samples but without ThT, were include for further TEM imaging and pFTAA end-point measurements.

Data were normalized and fitted in ThT kinetics Fitting formula

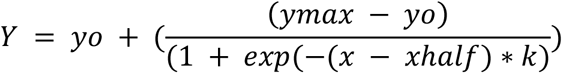

T_1/2_ and k was calculated from the formula above. Fluorescence amplitude was identified as the highest value of kinetics and lag time from, lagtime= t_1/2_-(2-k). Statistical analysis was performed using Brown-Forsythe and Welch ANOVA test with Dunnett T3 multiple comparisons correction and 95% confidence interval. Mean difference and 95%CI of difference was plotted. GraphPad was used for statistics and graphs.

### Transmission electron microscopy

Once the ThT signal reach a plateau, the resulting fibrils from the peptide:rAβ1-42 samples were analysed for their structural characteristics. Therefore, 10ul of each sample was spotted in a copper grid (Formvar/Carbon on 400 Mesh Copper - AGAR SCI, AGS162-4), previously glow discharged. The sample was adsorbed for 3 minutes. Afterwards the grids were washed by contact with three drops of MQ water, negative stained with one drop of uranyl acetate (2% w/v) for one minute, and finally washed in a drop of MQ water. The grids were examined using a JEM-1400 120kV transmission electron microscope (Jeol, Japan), at accelerating voltage of 80 keV. At least 9 positions on the grid was used for quantification. Fibrils were quantified if they were able to be traced from start to end. More than 100 fibrils were quantified in most cases (except rAβ1-42+P23, rAβ1-42+P24) (Appendix Table S7). Length was measured by using the freehand line of Fiji and tracing the fibrils from start to end. Statistical analysis was performed using Brown-Forsythe and Welch ANOVA test with Games-Howell multiple comparisons correction and 99% confidence intervals. GraphPad was used for statistics and graphs.

### Atomic force microscopy imaging

Fibril samples were deposited on freshly cleaved mica for AFM imaging. Each sample was adjusted using a solution of HCl at a predetermined concentration to result in the sample reaching pH 2. Immediately afterwards, 20 μl samples were deposited onto freshly cleaved mica surfaces (Agar scientific, F7013) and incubated for 5 minutes. Following incubation, the sample was washed with 1 ml of filter sterilised milli-Q water and then dried using a stream of nitrogen gas. Fibrils were imaged using a Multimode AFM with a Nanoscope V (Bruker) controller operating under peak-force tapping mode using ScanAsyst probes (silicon nitride triangular tip with tip height = 2.5-2.8 μm, nominal tip radius = 2 nm, nominal spring constant 0.4 N/m, Bruker). Each collected image was scanned at either 4 × 4 μm and 2048 × 2048 pixels or 8 × 8 μm and 4096 × 4096 pixels. Therefore, the same pixel density was maintained for all images within the dataset. A scan rate of 0.203 Hz was used. A noise threshold of 0.5 nm was used, and the Z limit was reduced to 1.5 μm. Nanoscope analysis software (Version 1.5, Bruker) were used to process the image data by flattening the height topology data to remove tilt and scanner bow. Fibrils were traced (Aubrey *et al.*, 2020; Xue *et al*, 2009), digitally straightened(Egelman, 1986), and surface envelope reconstructed as previously described(Aubrey *et al.*, 2020; Lutter *et al.*, 2020) using an in-house application. The height profile for each fibril was extracted from the centre contour line of the straightened fibrils from which the average height of each fibril was calculated.

### pFTAA and curcumin measurements

pFTAA and curcumin end-point measurements were performed on 3-day old peptide:Met-Aβ1-42 samples, prepared as described above, using a final pFTAA concentration of 0.5 μM and curcumin 5 μM. Measurements were done in a low-volume black 384-well plate (Corning) using a ClarioStar fluorescence plate reader (BMG Labtech). After excitation at 440 nM, the emission spectra of pFTAA or curcumin were measured between 468 nm and 650 nm. Curcumin spectra was normalized and tested for significance with Kolmogorov-Smirnov test using GraphPad. The ratio of the two pFTAA picks was calculated average(505,506,507)/average(530,539,540) and the statistical significance was calculated using Brown-Forsythe and Welch ANOVA test with Dunnett T3 multiple comparisons correction and 95% confidence interval using GraphPad.

### HEK Aβ1-42 biosensor cell line

The Aβ1-42 biosensor cell line was developed in house. Briefly, a Aβ1-42 gene block was cloned in the multiple cloning site of a lentivirus plasmid, containing mCherry tag at N-terminus and CMV as promoter. The plasmid was transfected in HEK cells, together with the packaging plasmids (pCMV-deltaR 8.9 and pCMV-VSV-G), for the production of viral particles. HEK cells were transduced with these viral particles and sorted for mCherry. Cells were diluted in 96 well plates and grown as single cell colonies. A colony showing diffused expression of mCherry-Aβ1-42 was expanded and stored, to further be used as a biosensor cell line in Aβ seeding assays.

### Seeding and transfection assay of biosensor Aβ1-42 cell line

The biosensor mCherry-Aβ1-42 HEK cell line was cultured in DMEM medium, supplemented with 10% FBS at 37°C, and a 5% CO_2_ atmosphere. The seeding assay was performed by transfecting these cells with freshly prepared rAβ1-42 seeds, described above, and the quantification of the formed Aβ1-42 inclusions, resulted from the Aβ1-42 aggregation.

Briefly, the assay was performed in 96-well plate (PerkinElmer), previously coated for 30’ with poly-L-lysine at 37°C and washed three times with PBS. Adhered cells were passed twice through a 22G needle and plated at 15.000 cells/well and 5h later were transfected with 0, 10, 50, 100nM seeds or 100ng of DNA per well, using Lipofectamine 3000 (Invitrogen) according to the manufacturer.

After 17 hours of seed transfection and 41h of DNA transfection, the cells were fixed with 4% formaldehyde in PBS for 10 minutes. Cells were washed with PBS, block and permeabilize with 1%BSA, 0.2% TritonX-100 in PBS for 1hour. Cells were nuclei stained with 3uM Draq7 in 1%BSA in PBS for 1h. Cells were washed and plates were imaged using Operetta CLS. For seeded cells: For each well 17 fields were imaged by using the channels Digital Phase Contrast, mCherry (Ex:530-560, Em:570-650), DRAQ7 (Ex:615-645, Em: 655-705). The images were analysed by Operetta CLS. Nuclei was detected with DRAQ7, Cytoplasm with Digital Phase Contrast. Spots measured on ROIs: Nuclei and Cell. For transfected cells: For each well 17 fields were imaged by using the channels Digital Phase Contrast, mCherry (Ex:530-560, Em:570-650), DRAQ7 (Ex:615-645, Em: 655-705), EGFP (Ex:460-490, Em: 500-550). The images were analysed by Operetta CLS. Nuclei was detected with DRAQ7, Cytoplasm with Digital Phase Contrast. Spots measured on ROIs: Nuclei and Cell. EGFP intensity was measured for each cell identified. The baseline EGFP was calculated in PBS treated cells. Every cell with higher fluorescence was identified as transfected with our plasmids. The number of spots were identified in cells with and without EGFP fluorescence. No of spots per cell was calculated from No of spots/ no of cells for EGFP positive and negative cells (Appendix Table S8). Statistical significance was calculated using Ordinary one-way ANOVA with Dunett multiple comparison correction and unpaired t-test for comparison between transfected/nontransfected cells. GraphPad was used for statistics and graphs.

### FRAP

Images were acquired on Nikon A1R Eclipse Ti confocal with Plan APO VC 60x oil lens. For m-cherry excitation we used 561,6nm laser line and emission was collected at 570-620nm. A pre-bleached image was acquired and ROI was bleached for 0.06,0.62,1.25sec at 100% power with 30 image acquisition for 1.91sec after each bleaching and 50 image acquisition after last bleaching. For seeds a similar protocol was used with bleaching for 0.06,0.62,1.25sec and 30 image acquisition of 0.97sec after each bleaching and 50 image acquisition after last bleaching. Live cell imaging was done at 37°C and CO_2_.

### Immunofluorescence

Cells were fixed with 4% formaldehyde in PBS for 10 minutes. Cells were washed with PBS, block and permeabilize with 1%BSA, 0.2% TritonX-100 in PBS for 1hour. Cells were stained with Alexa647 1:1000 in 1%BSA in PBS for 1h. Images were acquired on Nikon A1R Eclipse Ti confocal with Plan APO VC 60x oil lens. For m-cherry excitation we used 561,6nm laser line and emission was collected at 570-620nm and for Alexa647 we used a 640.8 laser and the emission was measured at 663-738.

### Proteomic analysis

Aβ sequence was divided into hexapeptides with a sliding window. Proteins with absent values, #Unique peptides<1 and Score Sequest HT<10 were filtered out from the proteomic dataset(Xiong *et al.*, 2019). The remaining proteins were searched for homology to Aβ sequence with up to 2 mutations using an inhouse algorithm. Proteins with a fold change>1.186 was previously characterized as upregulated in APs, and defined the AP proteins. Enrichment ratios were calculated by

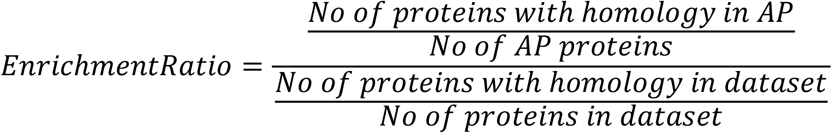

Significance was calculated with hypergeometric test with Bonferroni correction for multiple comparisons. A similar approach was used for analyzing nonAD proteomic data, mouse data and Glial Cytoplasmic Inclusions.

In order to verify the significance of the observed enrichments of homologues peptides in AD plaque proteins, we asked how (un)likely it would be to observe these enrichments if the assignment of “AD-plaque-enriched” proteins was entirely random. To do so, we generated 1000 random samples of the same size of the set of proteins found to be enriched in AD plaques by Xiong et al(Xiong *et al.*, 2019). For each of these random samples, we calculated logFCs for the homologues peptides versus background (either MS dataset background or the entire proteome, as indicated). We then calculated p-values for the observed logFCs versus the distribution obtained through random sampling, assuming normality. Same approach was used for nonAD and mouse plaques.

Aggregation propensity of hexapeptides with homology to overrepresented regions of Aβ from AP and non-AP proteins was calculated using TANGO. Statistical analysis was performed using Kolmogorov-Smirnov test. GraphPad Prism was used for statistical analysis.

Proteins identified in other MS studies were searched for homology to overrepresented Aβ APRs, and pie charts of the proteins with homology versus proteins with no homology were plotted using GraphPad Prism.

Gene Ontologies of AP proteins with and without homology to Aβ APRs were identified by using ClueGO(Bindea *et al*, 2009) plug-in of Cytoscape. Ontology used GO_BiologicalProcess-EBI-UniProt-GOA_18.09.2019. Statistical analysis was performed with Right-sided hypergeometric test with Bonferroni step down multiple correction. Identified pathways and their p-values were imported in REVIGO(Supek *et al*, 2011). Pathways were summarized using Homo Sapiens database and SimRel as semantic similarity measure. TreeMap R script was downloaded and the most significant pathway in the group was used as representation. Full GO in supplementary. A similar approach was used for GO identification of nonAD brain AP proteins.

Glial cytoplasmic inclusions analysis was performed as previously, using as AP proteins the ones identified in purified Lewy Bodies from at least 4 MSA cases (McCormack *et al.*, 2019). As background proteins we used proteins identified previously in human basal ganglia (Fernandez-Irigoyen *et al.*, 2014)

## Acknowledgements

The Switch Laboratory was supported by grants from the European Research Council under the European Union's Horizon 2020 Framework Programme ERC Grant agreement 647458 (MANGO) to JS, the Flanders institute for biotechnology (VIB), the University of Leuven (“Industrieel Onderzoeksfonds”), the Funds for Scientific Research Flanders (FWO) and the Flemish Agency for Work and Innovation (VLAIO). NL was supported by a post-doctoral fellowship from the FWO. LDA and WFX was supported by Biotechnology and Biological Sciences Research Council (BBSRC), UK grant BB/S003312/1.

LFR was supported by FWO 12N0319N and JdW by Alzheimer Research Foundation standard grant SAO-FRA 2019/0013 and Methusalem Grant of KU Leuven/Flemish Government The authors gratefully acknowledge Electron Microscopy Platform & Bio Imaging Core, Department of Neurosciences KU Leuven, VIB – KU Leuven Center for Brain & Disease Research for their support & assistance in this work. Nikon A1R Eclipse Ti confocal was acquired through a Hercules type 1 AKUL/097/037 grant to Wim Annaert.

---

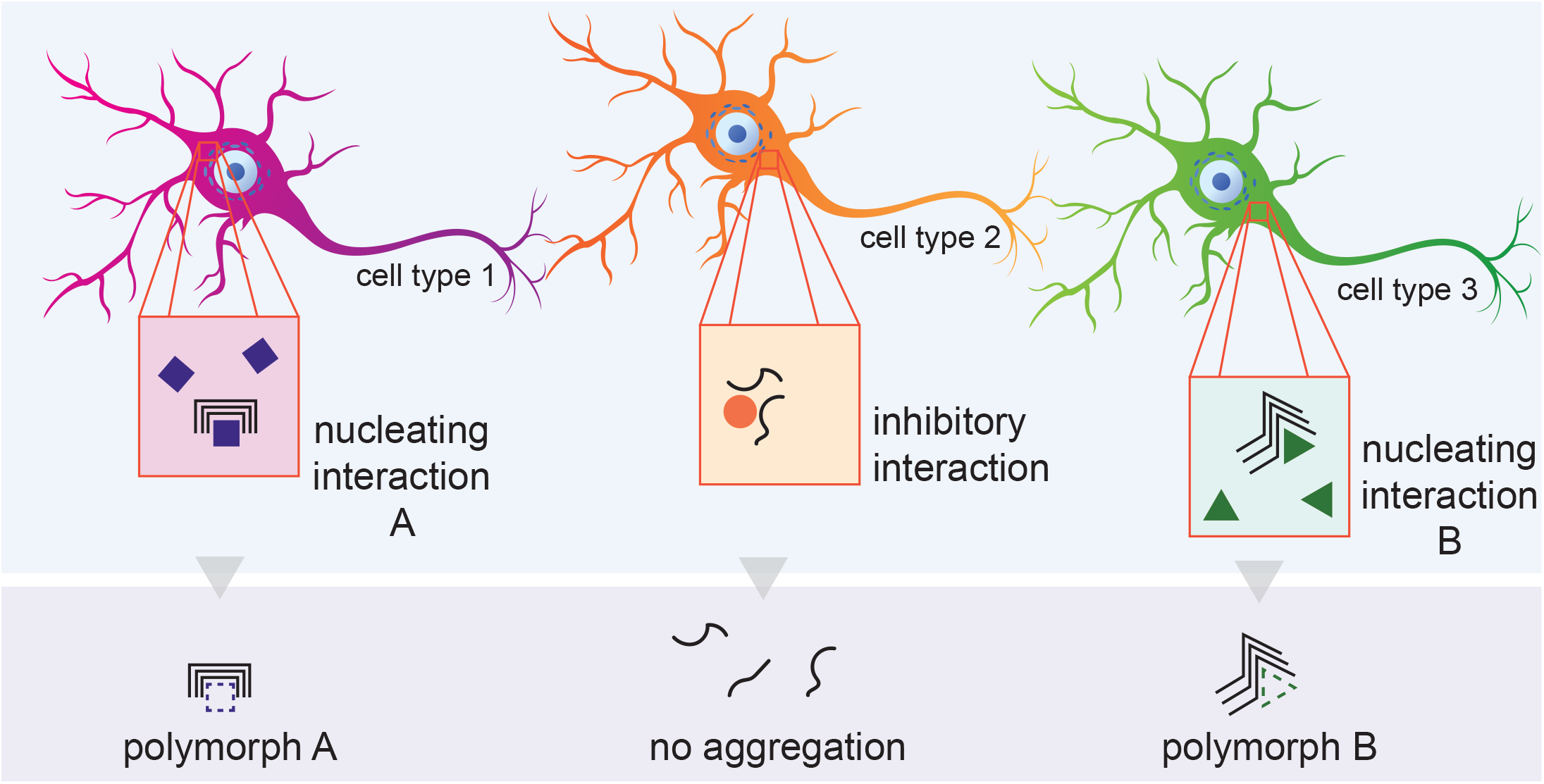

